# Hot water extract of *Arthrospira maxima* (AHWE) has broad-spectrum antiviral activity against RNA virus including coronavirus SARS-CoV2, and the antivirus spray application

**DOI:** 10.1101/2021.06.06.446935

**Authors:** Yi-Hsiang Chen, Ya-Chun Liao, Jyun-Yuan Huang, Yu-An Kung, Chuang-Chun Chiueh

## Abstract

The emergence and re-emergence of RNA virus outbreaks highlight the urgent need for the development of broad-spectrum antiviral agents. *Arthrospira maxima* has be used as a food source for a long time, and the protein or polysaccharide fractions were evidenced to have antiviral activity, therefore we examined the antiviral efficacy of hot water extract from *Arthrospira maxima* (AHWE), on Enterovirus 71 (EV71), Influenza virus, Herpes simplex virus (HSV), Respiratory syncytial virus (RSV), Ebola virus, and Coronavirus for antiviral spray application. In this study, we demonstrated that the AHWE shown 90 to 100% inhibition rate on the plaque formation of EV71, HSV-1, HSV-2, influenza virus, RSV, 229E and SARS-COV2 at virus attachment stage, and the long-lasting protection study also found while the AHWE was pre-exposed to the open air for more than 4 hours in plaque reduction assay. In addition, AHWE also had inhibitory effect on Ebola virus replication at 500 ug/ml. Finally, AHWE also shown no toxicity and skin sensitivity that imply it could be safe for future clinical use if approved by FDA. In conclusion, this study suggests that AHWE could be developed as a potential broad-spectrum antivirus spray product and therapeutic agent.

## INTRODUCTION

Many RNA viruses, such as Influenza virus, Enterovirus 71 (EV71), Severe Acute Respiratory Syndrome-Corona virus (SARS-CoV), Middle East respiratory syndrome coronavirus (MERS-CoV) and Zika virus had caused pandemics in the past 20 years. Recent emerging Severe Acute Respiratory Syndrome-Corona viruses 2 (SARS-CoV-2) cause outbreak of COVID19 still threats all mankind (Gao, 2018). In addition, Herpes simplex virus (HSV) and Respiratory syncytial virus (RSV) are also common pathogens that cause children and adults illness with no effective drug or vaccine for treatment and prevention. Vaccines and antiviral drugs usually target to specific virus species to prevent and control the spread of pathogens. However, vaccines and antiviral cannot be applied immediately to emerging viruses due to the viral specificity. Hence, antiviral drugs with broad-spectrum antiviral activity would be expected as effective approaches to control emerging and re-emerging viral infectious diseases. Although several broad-spectrum antiviral compounds are in preclinical studies or in clinical trials, to date, no drug has been approved (Ianevski et al., 2019).

Coronaviruse (CoV) is the largest known RNA viruse, composed of enveloped virus with positive single-stranded RNA. Its genome is contained in a capsid formed by nucleocapsid proteins included in an envelope. Three structural proteins are characteristic of coronavirus: membrane protein, envelope protein, and spike protein (S), a glycoprotein responsible for virus host cell attachment (Li, 2016). To date, seven coronaviruses have been found to cause diseases in humans (Zhu et al., 2020). While HKU1, NL63, OC43 and 229E are associated with mild symptoms in humans, SARS-CoV, MERS-CoV, and SARS-CoV-2, belonging to the betacoronavirus genus, cause severe to deadly pneumonia in humans (Corman et al., 2019).

The serious outbreak of the coronavirus disease of 2019 (COVID-19) caused by SARS-CoV-2 has created panic around the world because of its higher rate of infection. At the end of May, 2021, the confirmed cases reached 170 million along with 3.5 million deaths worldwide declared by World Health Organization (WHO). SARS-CoV2 uses ACE2 cellular receptor for entry into the host cell through binding of its spike (S) protein (Wan et al., 2020; Wu et al., 2020), leading to attack the lower respiratory system of the host and affect the lungs including pneumonia and failure of multiple organs (Jiang et al., 2020; Mallah et al., 2021). However, until now, there are still no effective drugs for the cure of individuals who are infected with the novel coronavirus, SARS-CoV-2.

*Arthrospira* is a genus of microscopic, multicellular, filamentous blue-green algae (cyanobacteria) and acts as a popular dietary supplement in human and as many animal species due to the rich in proteins, lipids, carbohydrate and elements. It is known as a rich source of proteins (55–70% of dry weight). It also contains lipids (5–6%), vitamins, minerals, and pigments (Hosseini et al., 2013). Many research studies show that *Arthrospira* has numerous health benefits, including antioxidant, immunomodulatory, anti-inflammatory, anticancer, anti-viral and anti-bacterial activities (Abdel-Daim et al., 2015; Chen et al., 2016; Hayashi et al., 1996; Wu et al., 2016).

Previous studies indicated that hot water extract (HWE) of *Arthrospira platensis* was found to inhibit the replication of Herpes simplex virus type 1 (HSV-1) in HeLa cells within the concentration range of 0.08-50 mg/ml in a dose-dependent manner. However, the HWE achieved a dose-dependent effect on viral penetration. At 1 mg/mL of the HWE was found to inhibit virus-specific protein synthesis without suppressing host cell protein synthesis if added to the cells 3 hr before infection. (Hayashi et al., 1993).

Calcium spirulan (Ca-SP), a sulfated polysaccharide isolated from hot water extract of *Arthrospira platensis*, selectively inhibited the penetration of virus into host cells. Retention of molecular conformation by chelation of calcium ion with sulfate groups was suggested to be indispensable to its antiviral effect. Ca-SP could inhibit the replication of several enveloped viruses, including HSV-1, human cytomegalovirus (HCMV), measles virus, mumps virus, influenza A virus, and human immunodeficiency virus (HIV-1)(Hayashi et al., 1996; Mader et al., 2016).

Chen et al. demonstrated that cold water extract of *Arthrospira platensis* has low cellular toxicity, and is well-tolerated in animal models at one dose as high as 5,000 mg/kg, or 3,000 mg/kg/day for 14 successive days. Anti-flu efficacy studies revealed that the *Arthrospira* extract inhibited viral plaque formation in a broad range of influenza viruses, including oseltamivir-resistant strains. Furthermore, *Arthrospira* extract was found to act at an early stage of infection to reduce virus yields in cells and improve survival in influenza-infected mice with inhibition of influenza hemagglutination (Chen, 2016 #12).

In this study, the hot water extract of *Arthrospira maxima* (AHWE) was evaluated for the antiviral efficacy on multiple viruses in vitro and in animal model.

## MATERIALS AND METHODS

### Cells

RD (Human Rhabdomyosarcoma), MDCK (Madin-Darby Canine Kidney) and Huh7 (human liver cell line) cells were grown in Dulbecco’s Modified Eagles’s Medium (DMEM) supplemented with 10% fetal bovine serum (FBS). RD cells were used for EV71 infection. MDCK cells were used for influenza A/WSN/33 virus infection. Huh7 cells were used for 229E coronavirus infection. HEp-2 (Human epithelial type 2), Vero and Vero-E6 (African green monkey kidney cell) cells were grown in Minimum Essential Medium (MEM) supplemented with 10% fetal bovine serum (FBS). HEp-2 cells were used for RSV infection. Vero cells were used for HSV-1 and HSV-2 infection. Vero-E6 cells were used for SARS-CoV-2 infection.

### Virus

Enterovirus 71/TW/4643/1998 was kindly provided from Jem-Ren Wang (National Cheng Kung University, Tainan, Taiwan). Influenza A/WSN/33(H1N1) was purchased from the American Type Culture Collection (ATCC). Respiratory syncytial virus (A2) was kindly receiving from Li-Min Huang (National Taiwan University Hospital, Taipei, Taiwan). HSV-1/3709/14, HSV-2/2934/14, Coronavirus 229E and SARS-CoV2 were provided from Chang Gung Memorial Hospital (Taipei, Taiwan). All work was performed with approved standard operating procedures and safety conditions in for SARS-CoV 2 in BSL-3 and other viruses in BSL-2.

### Hot water extract of *Arthrospira maxima*

Extracts were prepared a described elsewhere(Chen et al., 2016). Briefly, *Arthrospira maxima* powder provide from Far East Microalgae Ind. Co., Ltd. was suspended in distilled water, and treated with heat for 1∼3 hours. The supernatants were collected by centrifugation and lyophilized as AHWE powder for storage.

### Plaque reduction assay

Monolayer cells in 6-well plates (1× 10^6^ cells/well) were infected with the indicated viruses, with about 50 to 200 plaque-forming units (P.F.U.)/well considered by each virus plaque size. 500 ul of virus solution was first mixed with 500ul of AHWE (1 mg/mL) and incubated at room temperature for one hour. Then, the virus mixture was added to the cells in 6-well plates for absorption for one hour at room temperature. The cultures were subsequently overlaid with 3 mL of E0 medium containing 0.3% agarose, and then incubated at 37°C for 48 to 72 hours. Afterwards, cells were fixed with 10% formaldehyde for 1 hour, and then stained with 0.5 % of crystal violet for 15 minutes and washed out. For the experiment of pre-exposing AHWE to the air, the AHWE solution was added in an open dish that placed into hood for 4 to 24 hours. After the incubation, collected the exposed solution from the dish and added sterile water to refill the volume until reach the same volume to original solution. Then, this recovered solution was tested as proceed as above mentioned. Plaques in each well were visualized and counted to evaluate the anti-virus efficacy and the inhibition rate was calculated as follows: Inhibition rate= (reduced number of plaques in antiviral spray treated group/ number of plaques in virus-alone group) *100%.

### Primary plaque reduction assay for Ebola virus

Confluent or near-confluent cell culture monolayers of African green monkey kidney (Vero CCL81) cells in 12-well disposable cell culture plates were prepared. Cells were maintained in MEM or DMEM supplemented with 10% FBS. For antiviral assays the same medium is used but with FBS reduced to 2% or less and supplemented with 1% penicillin/streptomycin. The test compound mixs were prepared in 2X MEM or 2X DMEM and then combined with 2X agarose or 2X methylcellulose to equate to 10 ug/ml to 10000 ug/ml. The virus control and cell control will be run in parallel with each tested compound. Further, a known active drug is tested as a positive control drug (T-705 for Ebola Zaire) using the same experimental set-up as described for the virus and cell control.

The assay wass initiated by first removing growth media from the 12-well plates of cells, and infecting cells with approximately 100 plaque forming units (pfu) of virus. Cells were incubated for 60 min: 100 μl inoculum/ well, at 37°C, 5% CO2 with constant gentle rocking. Virus inoculum were removed, cells washed and overlaid with either 1% agarose or 1% methylcellulose diluted containing 2% FBS and 1% penicillin/streptomycin and corresponding AHWE concentration. Cells were incubated at 370C with 5% CO2 for 10 days. The overlay was then removed and plates stained with 0.05% crystal violet in 10% buffered formalin for approximately twenty minutes at room temperature. The plates were then washed, dried and the number of plaques manually counted. The number of plaques in each set of compound dilution was converted to a percentage relative to the untreated virus control. The 50% effective (EC50, virus-inhibitory) concentrations were then calculated by linear regression analysis (GraphPad).

Cell viability is evaluated in uninfected cell cultures in parallel to the actual primary PR assay. The cell viability assay (In vitro Toxicology Assay Kit, Neutral red (NR) based; Sigma) was being performed in 96-well plates following the manufacturer’s instructions. Briefly, growth medium was removed from confluent cell monolayers and replaced with fresh medium (total of 100µl) containing the test compound with the concentrations as indicated for the primary assay. Control wells contained medium with the positive control or medium devoid of compound. Wells without cells and growth medium only serve as blank. Typically, a total of three replicates will be performed for each condition. Plates were then incubated for 10 days at 37ºC with 5% CO2 according to the incubation time for Ebola plaque assays. The plates were then stained with 0.033% neutral red for approximately two hours at 37oC in a 5% CO2 incubator. The neutral red medium was removed by complete aspiration, and the cells rinsed 1X with phosphate buffered solution (PBS) to remove residual dye. The PBS was completely removed and the incorporated neutral red is eluted with 1% acetic acid/50% ethanol for at least 30 minutes. The dye content in each well is quantified using a spectrophotometer at 540 nm wavelength and 690 nm wavelength (background reading). The 50% cytotoxic (CC50, cell-inhibitory) concentrations were then calculated by linear regression analysis (GraphPad). The quotient of CC50 divided by EC50 gives the selectivity index (SI) value.

### Animal study

Far East Bio-Tec Co., Ltd. commissioned SGS Taiwan Ltd. to conduct all the animal study in this report, including acute oral toxicity study, skin irritation test and skin sensitization study. The studies were carried out in compliance with OECD Principles of Good Laboratory Practice (Organization for Economic Cooperation and Development, Paris, ENV/MC/CHEM (98) 17) and U.S. Food and Drug Administration Good Laboratory Practice Regulations, 21 CFR Part 58[Study Number: R-SR-KL20180408; Study Number: R-AOT-KL20180408; Study Number: R-GPMT-KL20180408]. Briefly, six of female SD rats were subjected into acute oral toxicity study. The rats were fasted overnight prior to dosing and were orally administered with single dose of AHWE in 2000 mg/kg body weight. Condition of animals were recorded at least once daily for total of 14 days. At the end of the test, animals were weighted and sacrificed for necropsies examination. For skin irritation test, furs of NZW rabbit backside from scapula to middle back were clipped to exposure skin surface. Sterile gauze was saturated with AHWE or distilled water and attached to the backside of NZW rabbit laterally for 4 hours. The dermal reactions were observed and recorded at 1, 24, 28, 72 hours after the treatment. The skin responses were checked and evaluated according to “Score System of Skin Reaction” described in Table 3. For skin sensitization study, the emulsion of AHWE solution (50 mg/mL in distilled water) or distilled water were mixed in 50% Freund’s complete adjuvant (volume ratio 1:1). Guinea pig backside were clipped and intradermally injected with 0.1 mL emulsions mentioned above. At 7-days post injection, gauzes statured with 0.5 mL AHWE solution were applied to injection site for 48 hours. At 21-day post injection, the patches soaked with 0.5 mL AHWE solution or distilled water were applied to the challenge site for 24 hours. The appearance of the challenge skin site of test and control group were observed at 24 and 48 hours after removal of patches. The skin reactions for erythema and edema were grading according to the “Magnusson and Kligman Scale” given in Table 5.

**Table 1.** The inhibition rate of AHWE against different viruses

| Plaque number | EV71 | FluA (H1N1) | FluA (H3N2) | FluB (Victoria) | RSV | HSV-1 | HSV-2 |
| --- | --- | --- | --- | --- | --- | --- | --- |
| Virus control | 97 | 60 | 63 | 82 | 105 | 200 | 50 |
| AHWE | 3 | 0 | 6 | 7 | 10 | 2 | 0 |
| Inhibition rate | 97 % | 100 % | 90.4 % | 91.4 % | 90.4% | 99 % | 100 % |

**Table 2.** The inhibition rate of AHWE exposed to the air more than 4 hours against different viruses.

| Plaque number | EV71 | FluA (H1N1) | FluA (H3N2) | FluB (Victoria) | RSV | HSV-1 | HSV-2 |
| --- | --- | --- | --- | --- | --- | --- | --- |
| Virus control | 97 | 60 | 105 | 82 | 50 | 97 | 60 |
| AHWE | 4 | 4 | 12 | 3 | 0 | 4 | 4 |
| Inhibition rate | 95.8% | 93.3% | 88.6% | 98.5% | 100% | 95.8% | 93.3% |

**Table 3.** The inhibition rate of AHWE against Ebola Zaire virus.

| Drug name | EC50 | CC50 | SI50 |
| --- | --- | --- | --- |
| Favipiravir | 96 ug/ml | >1000 ug/ml | >10 |
| AHWE | 500 ug/ml | >10000 ug/ml | >20 |

EC50 : compound concentration that reduces viral replication by 50%
CC50: compound concentration that reduces cell viability by 50%
SI50:CC50/EC50

**Table 4.** Score system of skin reaction in skin irritation test

| **Reaction** | **Primary irritation Score** |
| --- | --- |
| **Erythema and eschar formation** |  |
| ● No erythema | 0 |
| ● Very slight erythema (barely perceptible) | 1 |
| ● Well-defined erythema | 2 |
| ● Moderate erythema | 3 |
| ● Severe erythema (beet redness) to eschar formation preventing grading of erythema | 4 |
| **Oedema formation** |  |
| ● No oedema | 0 |
| ● Very slight oedema (barely perceptible) | 1 |
| ● Well-defined oedema | 2 |
| ● (edges of area well-defined by definite raising) |  |
| ● Moderate oedema (raised approximately 1 mm) | 3 |
| ● Severe oedema | 4 |
| ● (raised more than 1 mm and extending beyond exposure area) |  |

**Table 5.** Magnusson and Kligman scale for scoring the response in skin sensitization study

| **Reaction** | **Grading scale** |
| --- | --- |
| No visible change | 0 |
| Discrete or patchy erythema | 1 |
| Moderate and confluent erythema | 2 |
| Intense erythema and swelling | 3 |

## Results

### Broad spectrum antivirus activity of AHWE

The anti-viral efficacy of AHWE on EV71, Influenza A, RSV, HSV-1 and HSV-2 were evaluated by using plaque reduction assay. Each virus was mixed with AHWE, respectively, and then incubated with cells for viral infection. The final results of plaques formation were shown in Figure 1 and Figure 2. The inhibition rates of AHWE on EV71, Influenza A, RSV, HSV-1 and HSV-2 were summarized in Table.1. The efficacy of AHWE reduced the plaques formation is 97% (94/97) on EV71, 99% (198/200) on HSV-1, 100% (50/50) on HSV-2, 100% (60/60) on influenza A/WSN/33 virus and 90.4% (95/105) on RSV/A2. For long-lasting protection activity evaluation, the AHWE was pre-exposed to the air-flow in laminar flow for more than 4 hrs, and then recovered solution was tested in plaque reduction assay. The pre-exposed AHWE still inhibited 95.8% (93/97) of EV71 plaque formation. In addition, pre-exposed AHWE also inhibited 98.5% (197/200) of HSV-1, 100% (50/50) of HSV-2, 93.3% (56/60) of influenza A/WSN/33 and 88.5% (93/105) of RSV/A2 virus plaque formation (Table 2). These results revealed the AHWE obviously inhibited the replication of different virus at attachment stage of virus infection. The anti-viral activity of AHWE remained even after the exposure to the air for 4 hours.

**Figure 1.**
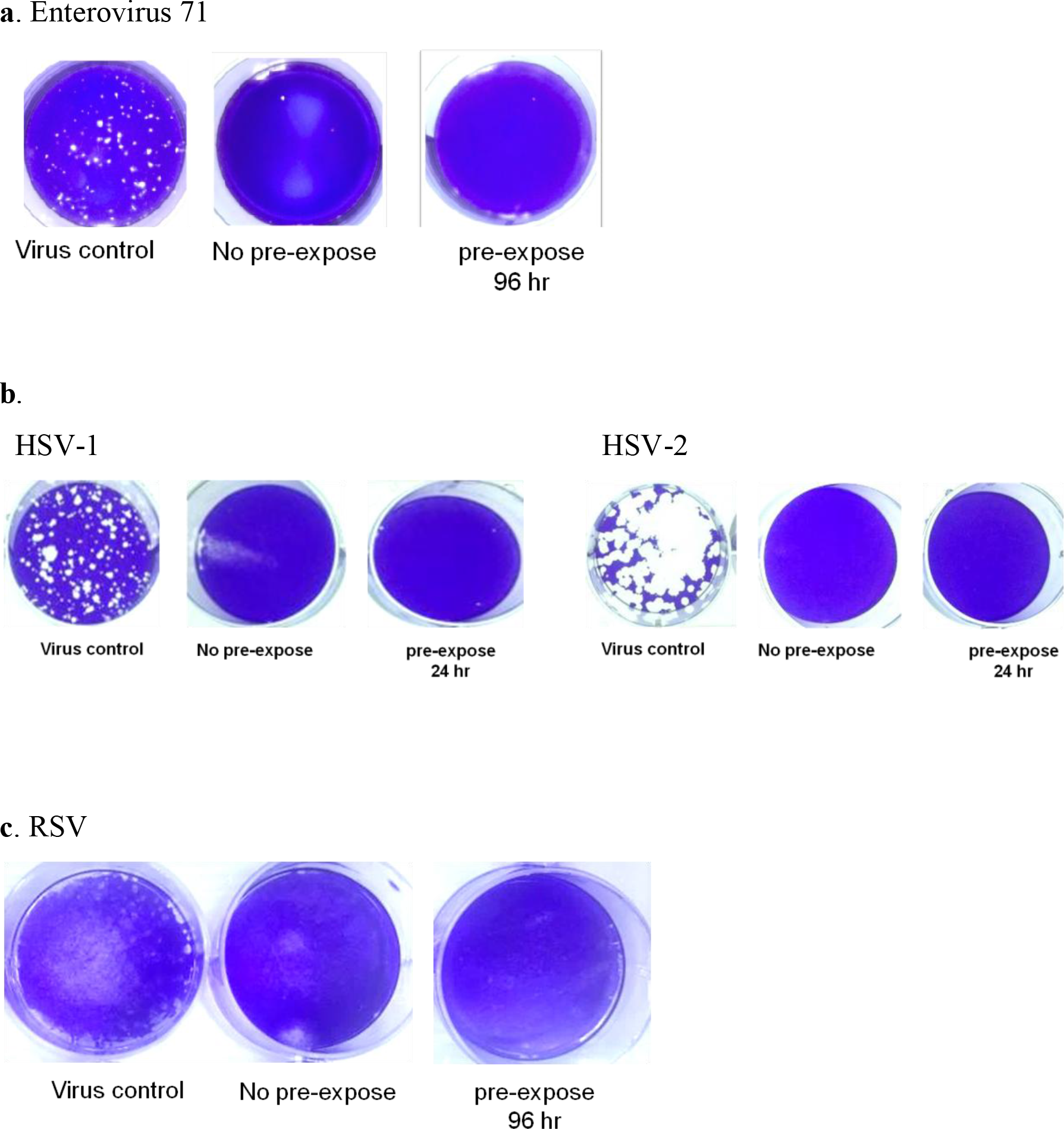
Antiviral activity of AHWE against various viruses by plaque reduction assay. Each kind of virus was mixed with AHWE, respectively, and then incubated with cells for viral infection. The viral yields were determined by the plaque assay, and represented as the inhibition rate (%). Representative plaques of the antiviral evaluation of AHWE against EV71 (a), HSV-1 and HSV-2 (b), and RSV(c) on cells were shown.

**Figure 2.**
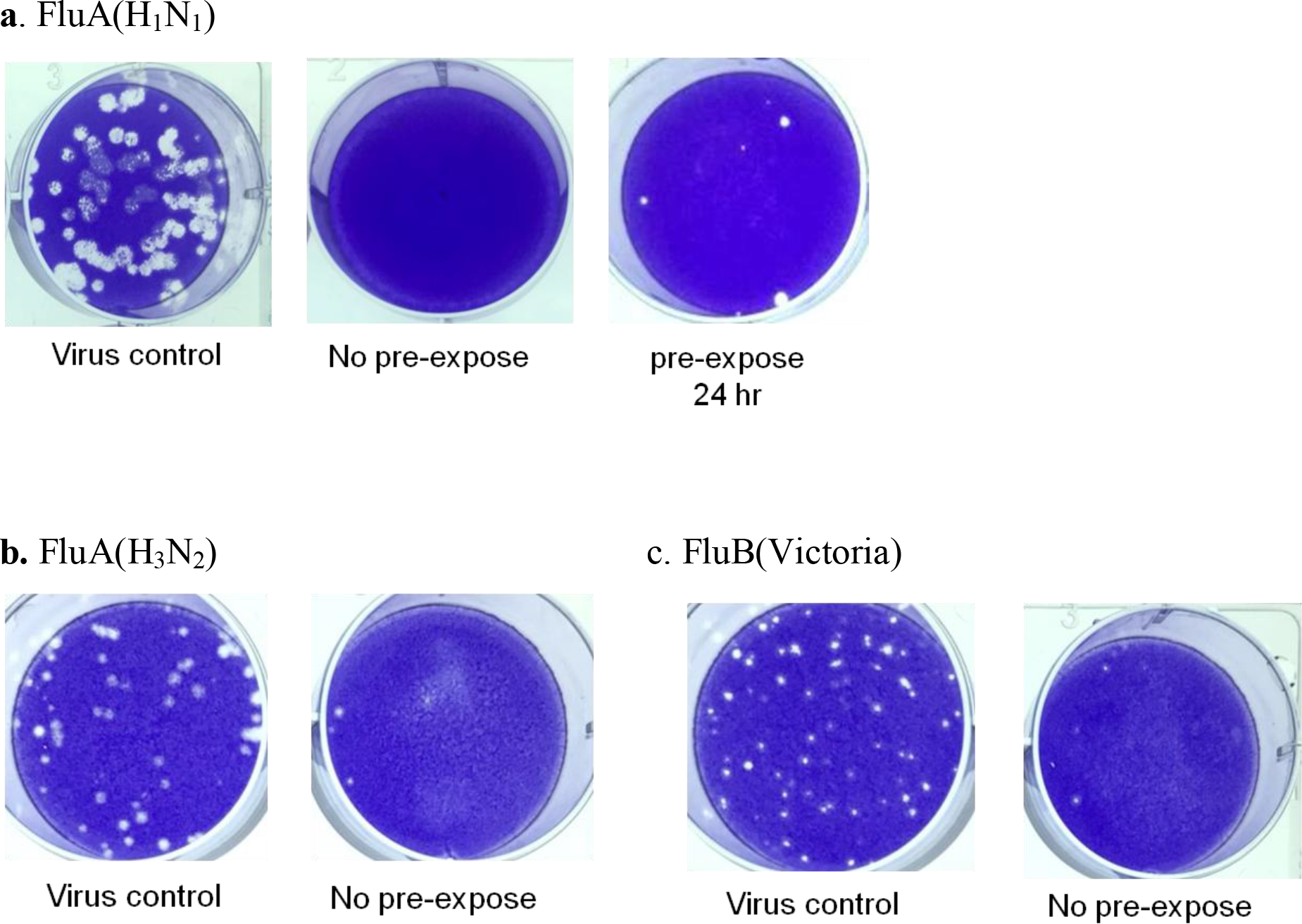
Antiviral activity of AHWE against influenza virus by plaque reduction assay. Influenza virus was mixed with AHWE and then incubated with MDCK cells for viral infection. The inhibition rate were determined by the plaque assay, and represented as the inhibition rate (%). The figure represents representative plaques of the antiviral evaluation of AHWE against Influenza virus on MDCK cells.

To explore the potential effect of AHWE on highly lethal viruses, we also utilized the non-clinical and pre-clinical services program offered by the National Institute of Allergy and Infectious Diseases to screen several highly lethal viruses, such as Ebola virus. The primary plaque reduction assay shown that AHWE inhibited the plaque formation of Ebola virus in Vero cells at 500 ug/ml without any cellular toxicity (Table 3). This result indicated that AHWE also had inhibitory effect on Ebola virus replication.

### The anti-Coronavirus on 229E and SARS-CoV2 activity of AHWE even pre-exposed to the air for more than four hours

Since the outbreak of COVID-19 caused by SARS-CoV-2 still threat all the world, the antivirus activity on coronavirus was critical for disease control. The efficacy of inhibition was also evaluated in plaque reduction assay that Huh7 cells were infected with 229E and VeroE6 cells were infected with SARS-CoV-2, respectively. The efficacy of AHWE reduced the plaques formation efficacy is 95% (1/20) on 229E coronavirus (Figure 3a) and 94.7% (6/113) on SARS-CoV-2(Figure 3b). For long-lasting protection activity evaluation, AHWE inhibited the 100% (0/20) of 229E and 95.6±2.35 of SARS-CoV-2 plaque formation while AHWE was pre-exposed to air for 4 hours (Figure 3).

**Figure 3.**
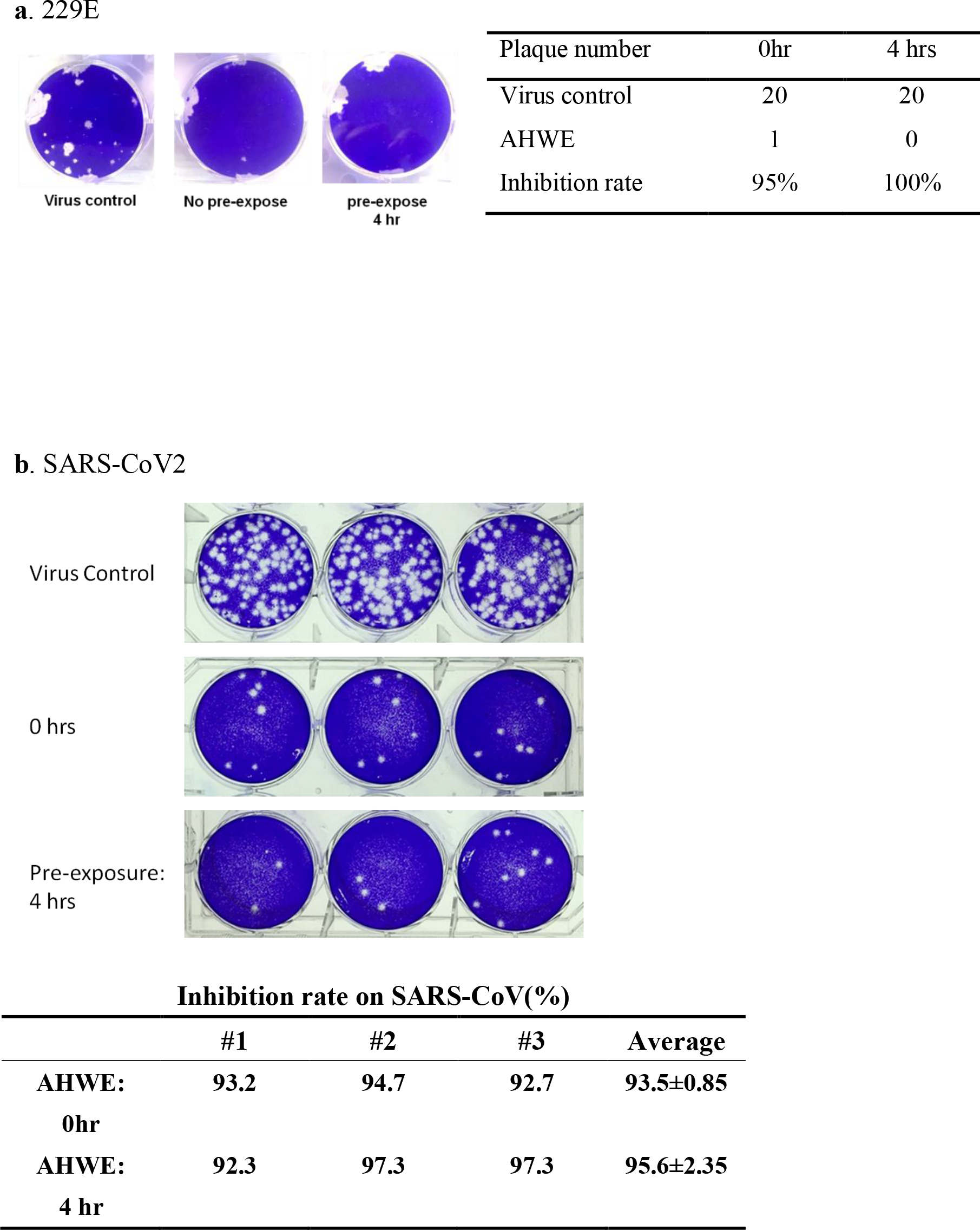
Inhibition rate of AHWE against coronavirus. The figure represents representative plaques of the antiviral evaluation of AHWE against 229E coronavirus (a) and SRARS-CoV-2(b) on cells. The table represents inhibition rate (%).

### AHWE is a potential anti-viral agent without significant toxicity in animal model

To explore the potent usage of AHWE, acute oral toxicity of AHWE was been examined on animal. SD rats were orally administrated with AHWE (2000 mg/kg body weight) and the animals were following up to 2 weeks for observing any clinical signs occurred. The body weight of rats had no obviously changed within 15 days post AHWE administration (Figure 4.) There were no animals died and no adverse effects were observed during experiment. In addition, no abnormal pathological sign was found in the necropsy at day 15 after AHWE administration. Therefore, the median lethal dose (LD_50_) of AWHE was estimated to be higher than 2000 mg/kg in the rats, which is considered to be safe.

**Figure 4.**
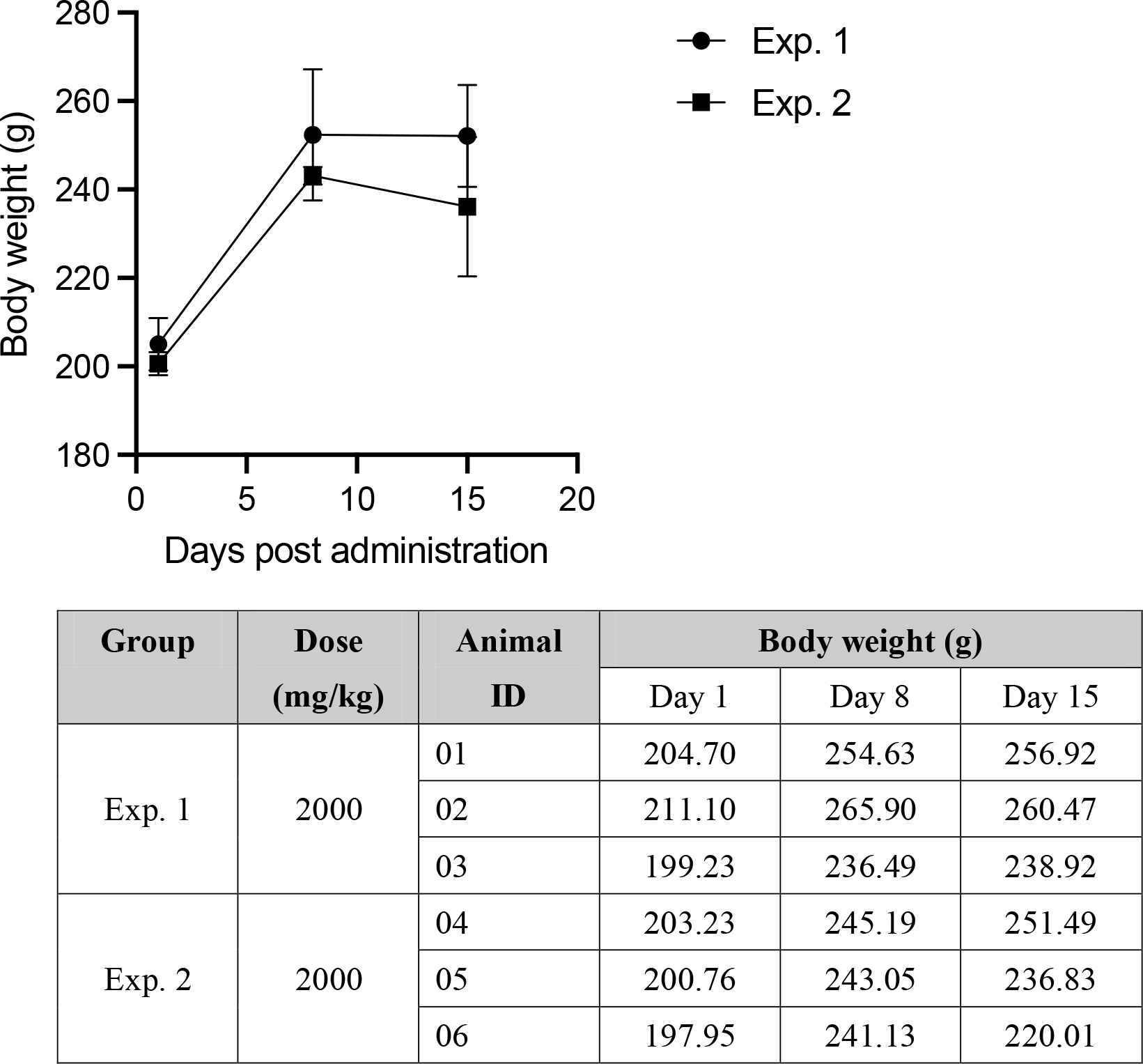
The body weights of SD rats were not significantly changed after AHWE administration orally. Six SD rats were orally administrated with AHWE solution (2000 mg/kg body weight) and their body weights were monitored at day 1, day 8 and day 15 post treatment.

Furthermore, to assess whether AHWE is suitable for epidermal usage, we conducted skin irritation test and skin sensitization study (maximization test) to avoid potential physiological damage on the skin. For skin irritation test, backsides of NZW rabbits were covered with two independent gauzes laterally, which saturated with 0.5 mL AHWE (50 mg/mL in distilled water) or distilled water. After 4 hours treatment, the dermal reactions at the treated areas were observed for up to 72 hours. The observation endpoints included erythema, edema, and any other toxicity responses. The score system of skin reaction was shown in Table 4. No obvious edema was found in either AHWE group or distilled water group. However, slight erythema was observed in the AHWE group (Figure 5). The results demonstrated that AHWE were regard as safe but might causing slight erythema with single topical application.

**Figure 5.**
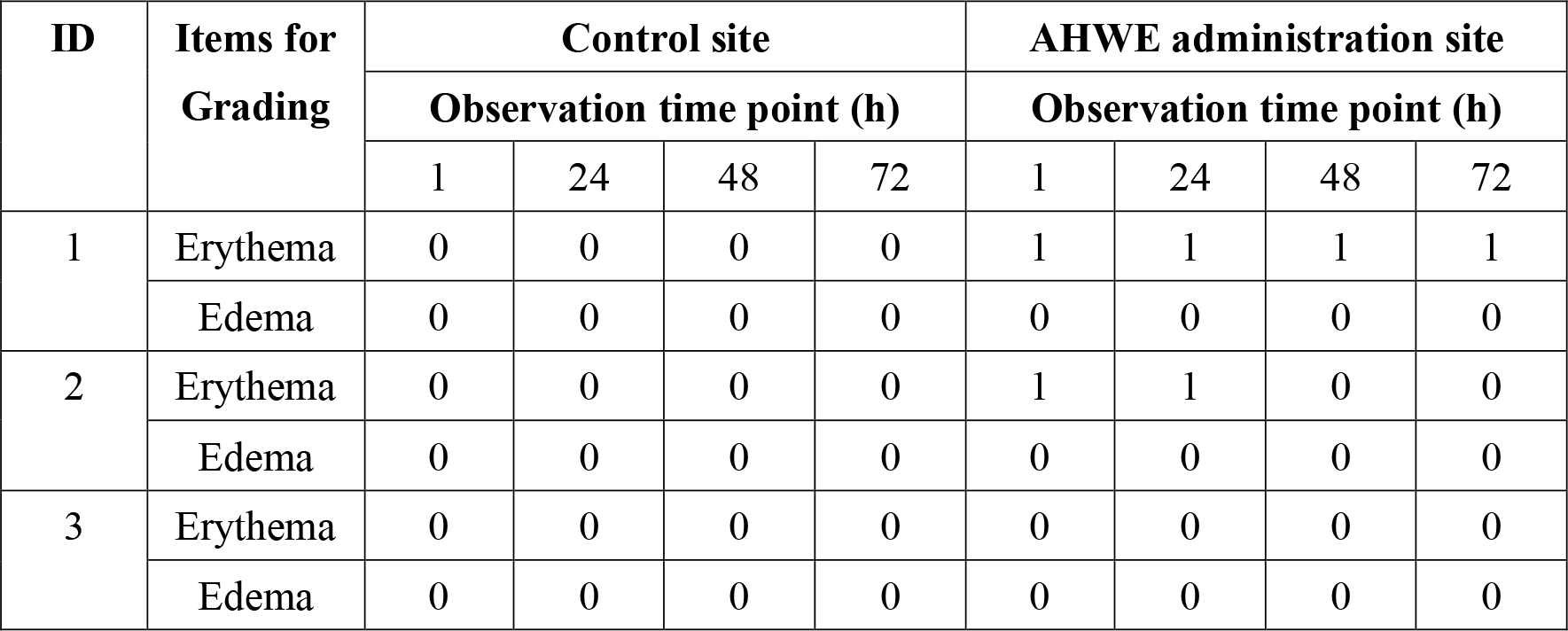
Administration of AHWE has no significant skin response in skin irritation test. NZW rabbits were treated with gauze for 4 hours at the backside, which soaked with AHWE or distilled water, and treated areas were observed as indicated time point after removal of gauze. The responses of skin were scored according to “Score System of Skin Reaction” described in Table 4.

For skin sensitization study, backsides of guinea pig were intradermally injected with AHWE (2.5 mg in emulsion of distilled water and Freund’s complete adjuvant) and the appearances of skin were observed after re-challenging. Skin reactions for erythema and edema were graded according to the “Magnusson and Kligman Scale” given in Table 5. As shown in Figure 6, the results indicated that AWEH did not produced skin sensitization in guinea pigs. In summary, AWEH was a potential broad-spectrum antiviral agent and safe for topical usage.

**Figure 6.**
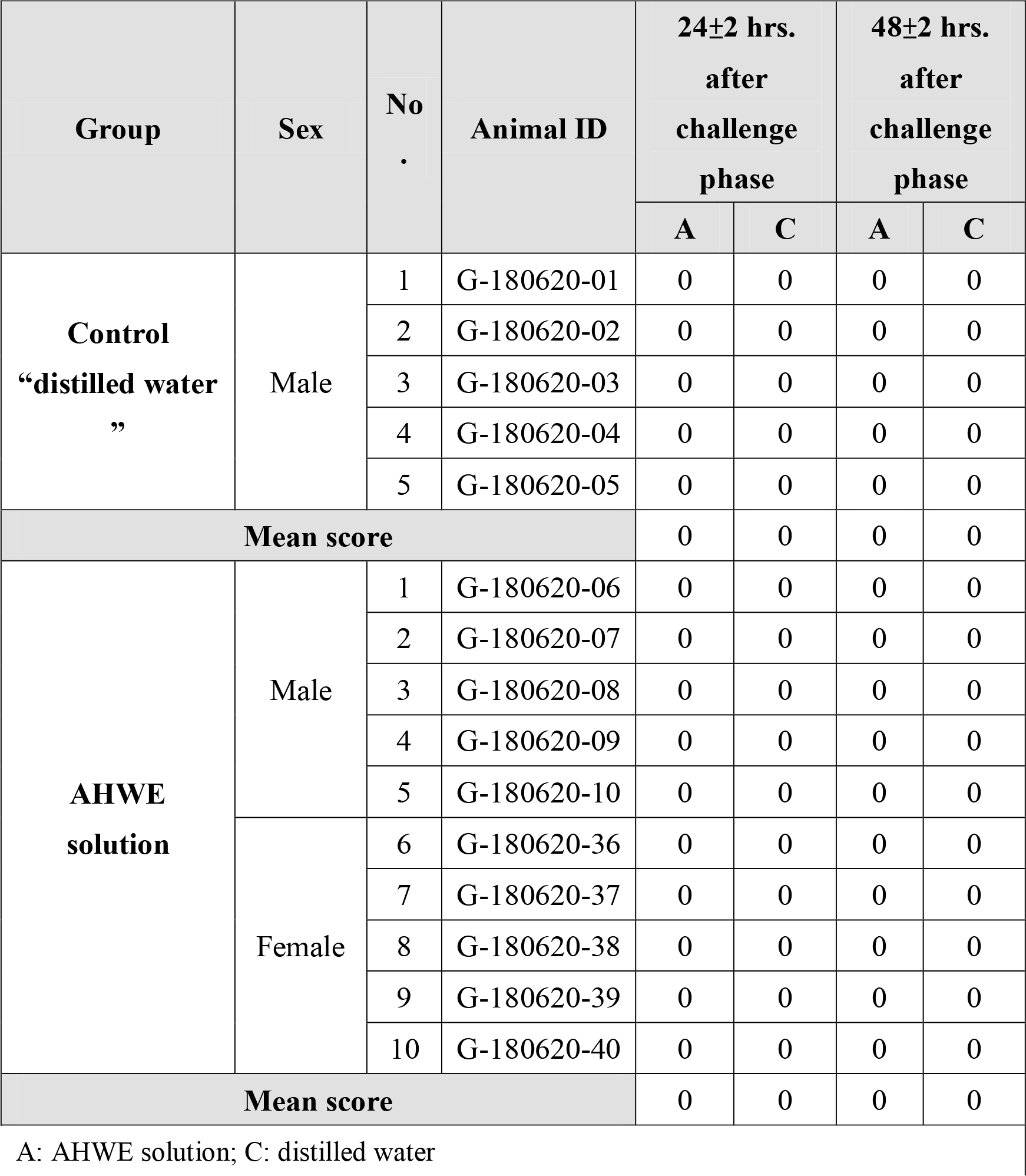
Administration of AHWE did not induce obvious response in skin sensitization study. Guinea pigs were intradermally injection with AHWE solution or distilled water at the backside and were re-stimulated by gauze soaked with AHWE solution after 7-days post injection. At 21-days post injection, treated site were re-challenged with the patches that soaked with AHWE solution or distilled water for 24 hours. The appearance of challenge sites were observed at indicated time points and grading according to the “Magnusson and Kligman Scale” given in Table 5.

## Discussion

The emergence and re-emergence of RNA virus outbreaks in past 20 years highlight the urgent need for the development of broad-spectrum antiviral agents. Most of these emerging viruses are highly pathogenic with limited therapeutic strategies. In this report, we demonstrated that the AHWE had been found to obviously inhibit the EV71, HSV-1, HSV-2, influenza A/WSN/33 virus, RSV/A2, 229E and SARS-COV2 plaque formation at virus attachment stage, and the long-lasting protection study also found even the AHWE pre-exposed to the open air for more than 4 hours in plaque reduction assay. Moreover, AHWE also had inhibitory effect on the replication of Ebola virus. Furthermore, the safety tests including oral acute toxicity, skin irritation evaluation and skin sensitivity evaluation had been performed and show no safety concern. The result reveals the ability of AHWE to prevent the contact transmission of broad-spectrum virus included the SARS-CoV-2.

The inhibition of viral entry is a key target for the development of antiviral agents. Some medical plants and natural compounds have been reported to have a wide range of antiviral activities. Among these, *Arthrospira* extracts have been reported to inhibit attachment of enveloped viruses such as HCMV, HIV, and Influenza virus. Although, the active components in AHWE is not identified in this study, but the mechanism of action of *Arthrospira* extract against these tested viruses may be similar. The active components in AHWE might bind to enveloped glycoproteins of SARS-CoV-2 and play a role in the inhibition of virus entry. In summary, we have successfully identified the broad-spectrum of antiviral activity of AHWE on EV71, HSV, Influenza, RSV, 229E and SARS-COV2 that could be a promising antiviral agent for emerging viruses.

The chemical characterization of AHWE includes 42% of polysaccharide, 6% of protein, 20% of nucleic acid, and 11% of ash. The heat, acid, and basic-stable characterization of AHWE also would be applied as a long-lasting hand sanitizer to prevent the virus transmission across the environment.

## Acknowledgments

We thank the technology support from Research Center for Emerging Viral Infections, Chang Gung University, Taiwan for virus infection and analysis. We also thank the non-clinical and pre-clinical services program offered by the National Institute of Allergy and Infectious Diseases (NECA#2014-001-01).

## Notes

### Competing Interest Statement

The authors have declared no competing interest.

